# A large nuclear and mitochondrial sequence dataset for phylogenetic analysis of the hyperdiverse Asterophryinae frogs of the New Guinea region with data on lifestyle, GPS coordinates, elevation, and multispecies communities with accompanying code

**DOI:** 10.1101/2022.09.15.508180

**Authors:** Ethan C. Hill, Mary J. Jarman, Claire J. Fraser, Diana F. Gao, Elizabeth R. Henry, Allison R. Fisher, Bulisa Iova, Allen Allison, Marguerite A. Butler

## Abstract

The data provided here is related to the article “Resolving the Deep Phylogeny: Implications for Early Adaptive Radiation, Cryptic, and Present-day Ecological Diversity of Papuan Microhylid Frogs” [1]. The dataset is based on 233 tissue samples of the subfamily Asteroprhyinae, with representatives from all recognized genera, in addition to three outgroup taxa. The sequence dataset contains over 2400 characters per sample for five genes (three nuclear loci: BDNF, NXC-1, and SIA; and two mitochondrial loci: ND4 and CYTB) and is 99% complete. New primers were designed for all loci and accession numbers for the raw sequence data are provided. The sequences are used with geological time calibrations to produce time-calibrated Bayesian inference and Maximum Likelihood phylogenetic reconstructions using BEAST2 and IQTREE. Lifestyle data (arboreal, scansorial, terrestrial, fossorial, semi-aquatic) were collected from the literature and field notes and used to infer ancestral character states for each lineage. Collection location data (GPS and elevation data) were used to verify sites where multiple species or candidate species co-occur. All sequence data, alignments, and associated metadata (voucher specimen number, species identification, type locality status, GPS, elevation, site with species list, and lifestyle) as well as the code to produce all analyses and figures are provided.

## Specifications table

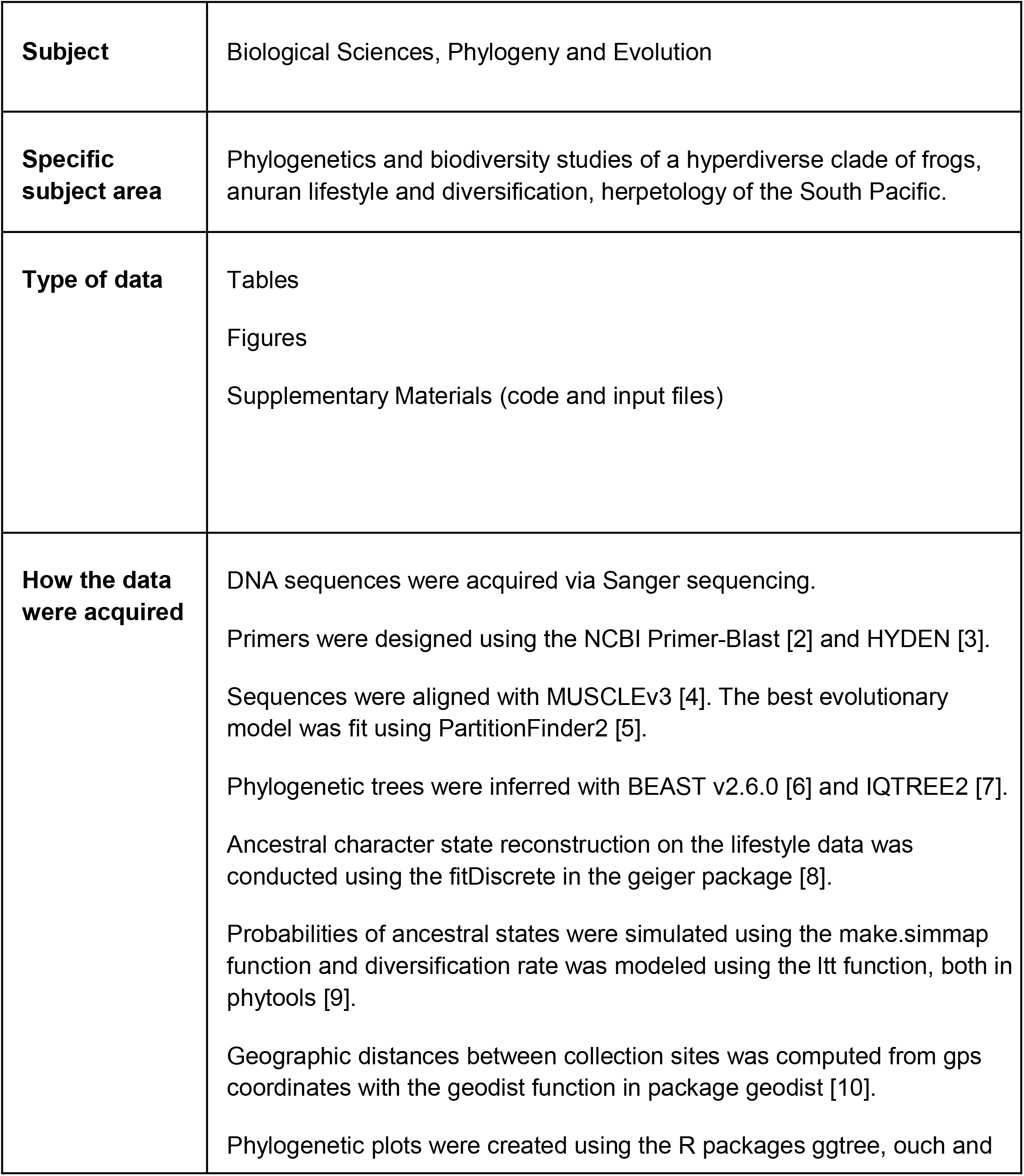
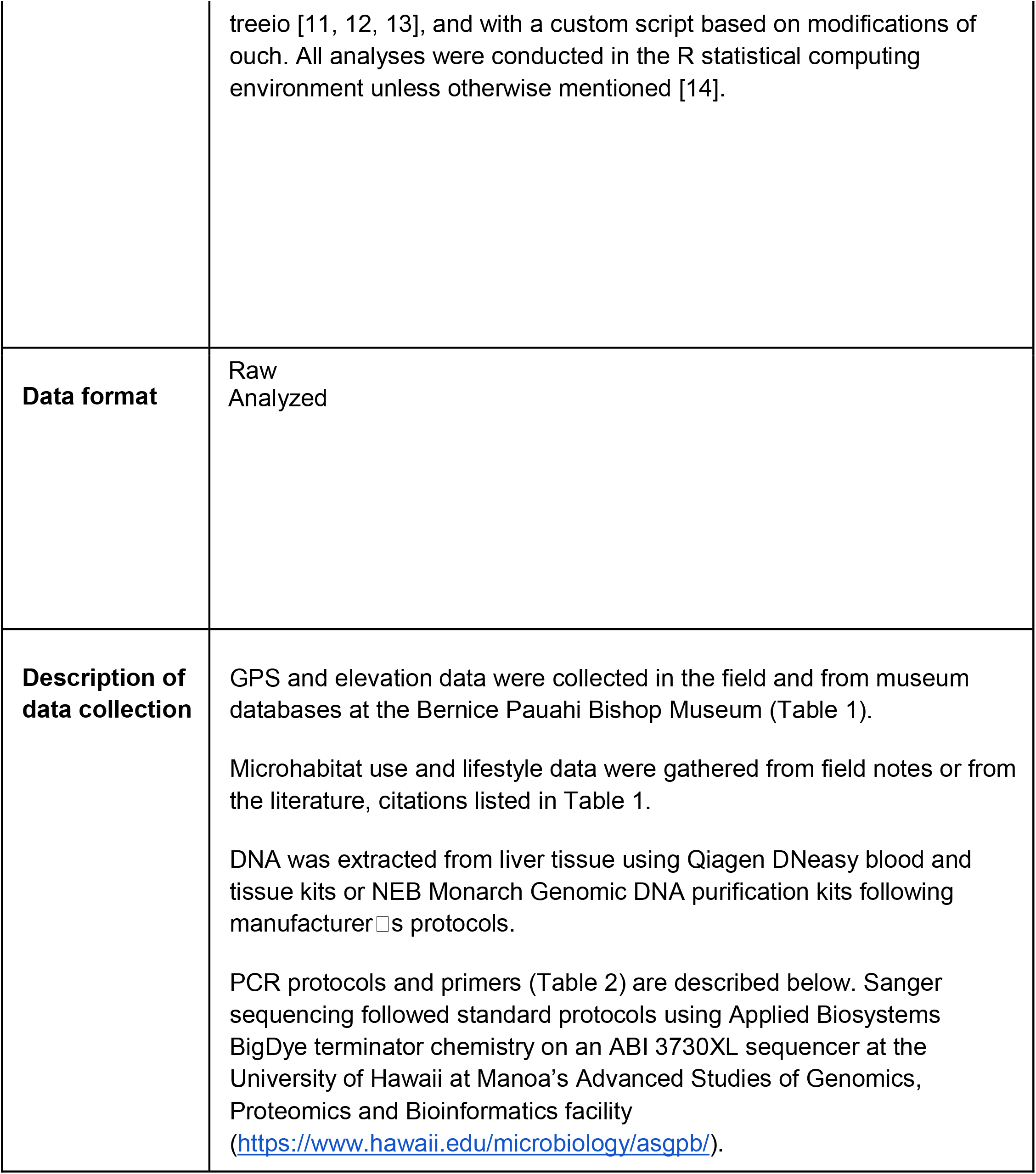
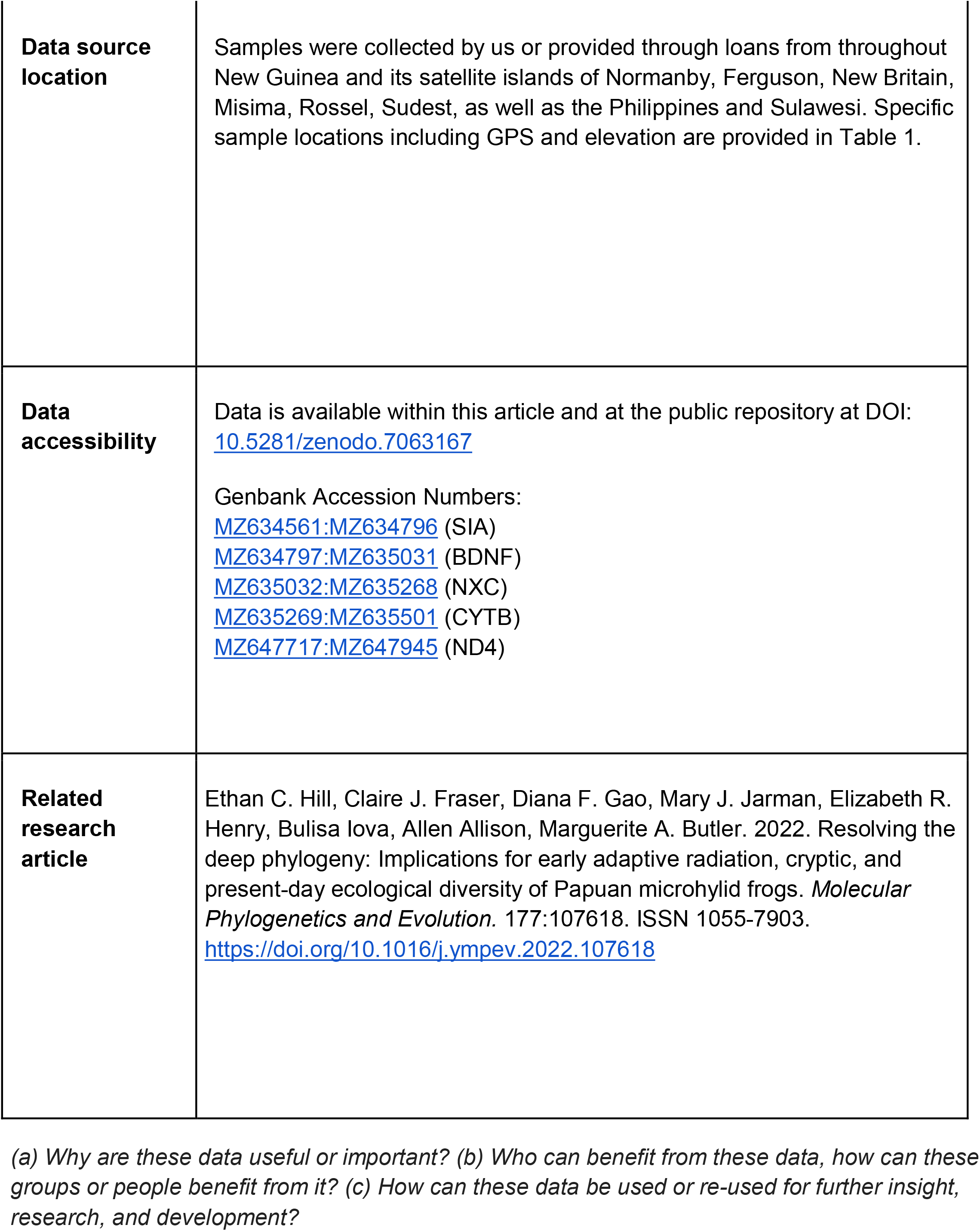

## Value of the data

- Molecular phylogenetic studies of the hyperdiverse Asterophryinae frogs have proved insufficient for resolving the deep nodes of this radiation. The nuclear and mitochondrial Sanger sequence data presented here for 205 species, combined with large geographic sampling and a 99% complete sequence dataset produce a nearly completely resolved and strongly supported phylogeny, clarifying intergeneric relationships, identifying many cryptic species complexes, and resolving many taxonomic questions.
- The phylogenetic data is complemented by associated GPS and lifestyle data from 77 sites across all major geological sectors of New Guinea and several satellite islands. The dataset includes complete species sampling in 18 multi-species communities ranging from two to 15 species.
- New primers are presented for three nuclear and five mitochondrial loci which resulted in 99% completeness of the molecular dataset, and will be useful for sequencing other vertebrates (nuclear loci) or anurans (mitochondrial loci).
- These data will be foundational for future studies of biogeography, pacific biodiversity, ecological diversification, community assembly, character evolution, and exploring the tempo and mode of evolution. This dataset will also be useful for exploring the impact of data completeness for sequence or geographical sampling, and rigorous model fitting on resolving large phylogenies for clades that have undergone rapid diversification.

## Data description

Asterophryinae is a hyperdiverse subfamily of microhylid frogs that has remained poorly resolved despite decades of phylogenetic work [reviewed in 1]. Given the ∼20 MY age of the clade and the poor resolution of deeper nodes in previous efforts, we suspected that long-branch attraction was a problem preventing resolution of nodes along the backbone of the phylogeny, and that mitochondrial loci alone were not sufficient to clarify intergeneric relationships. In addition, previous studies included only a small fraction of the clade□s diversity. In this study, we present a well resolved phylogeny for 205 species (233 taxa) of Papuan Asterophyrinae using both Bayesian inference (Figure 1) or Maximum Likelihood (Figure 2) reconstruction methods, that importantly expands geographic sampling and tests whether widely-distributed species instead represent cryptic diversity. These trees show well-defined genera and clarified the majority of intergeneric relationships, and also clarified cryptic species complexes for many taxa that were once believed to be widespread. Instead, the data reveal a mosaic of species with small geographic ranges and a tendency toward fragmentation. Geographic distance between tips in the tree can be obtained from GPS coordinates (Table 1), and used in analyses of geographic range evolution.

**Figure 1.**
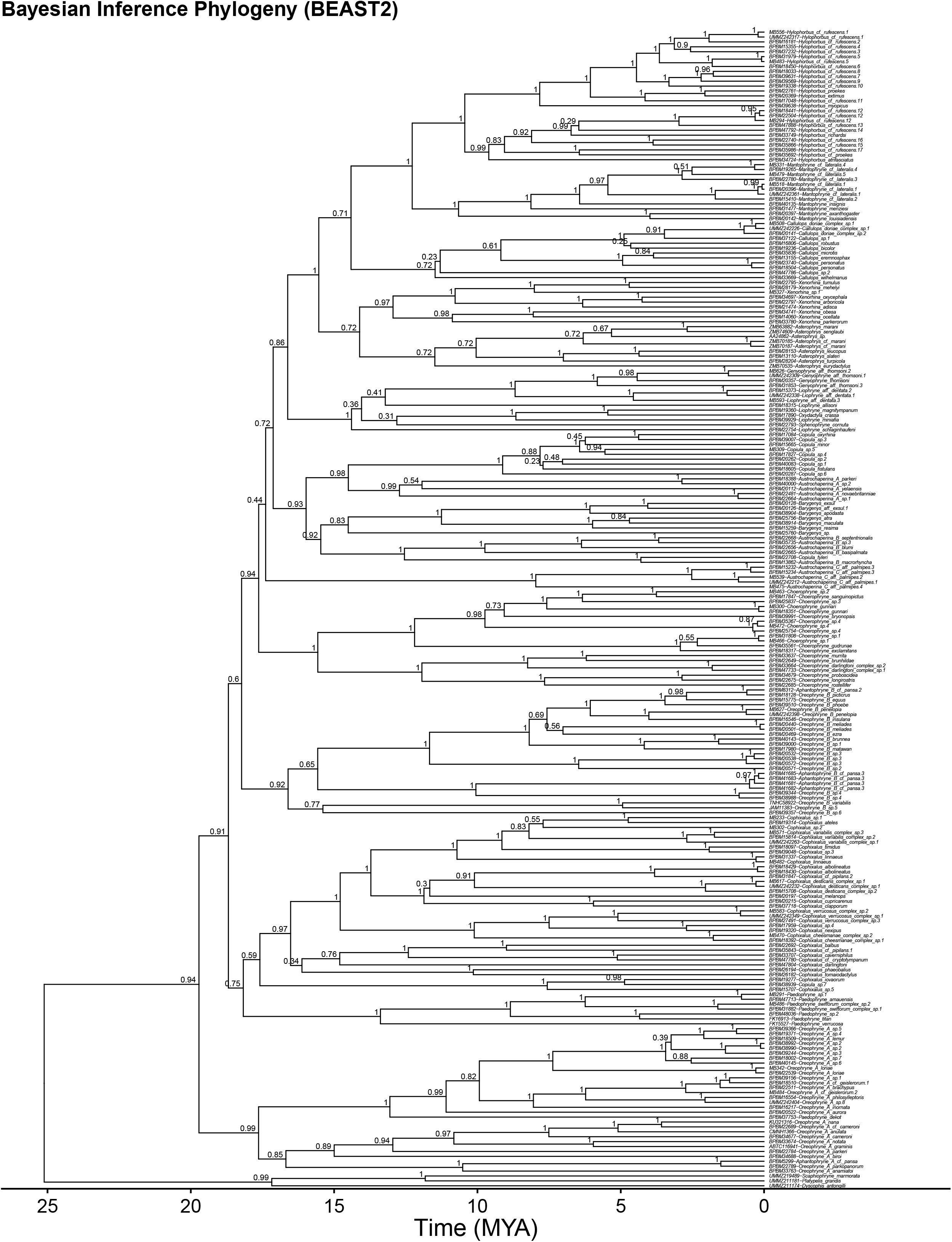
Time calibrated Bayesian inference phylogeny for 233 samples of Asterophryinae and three outgroups generated using BEAST2. Posterior probabilities are provided above nodes. All trees contain the same set of tips.

**Figure 2:**
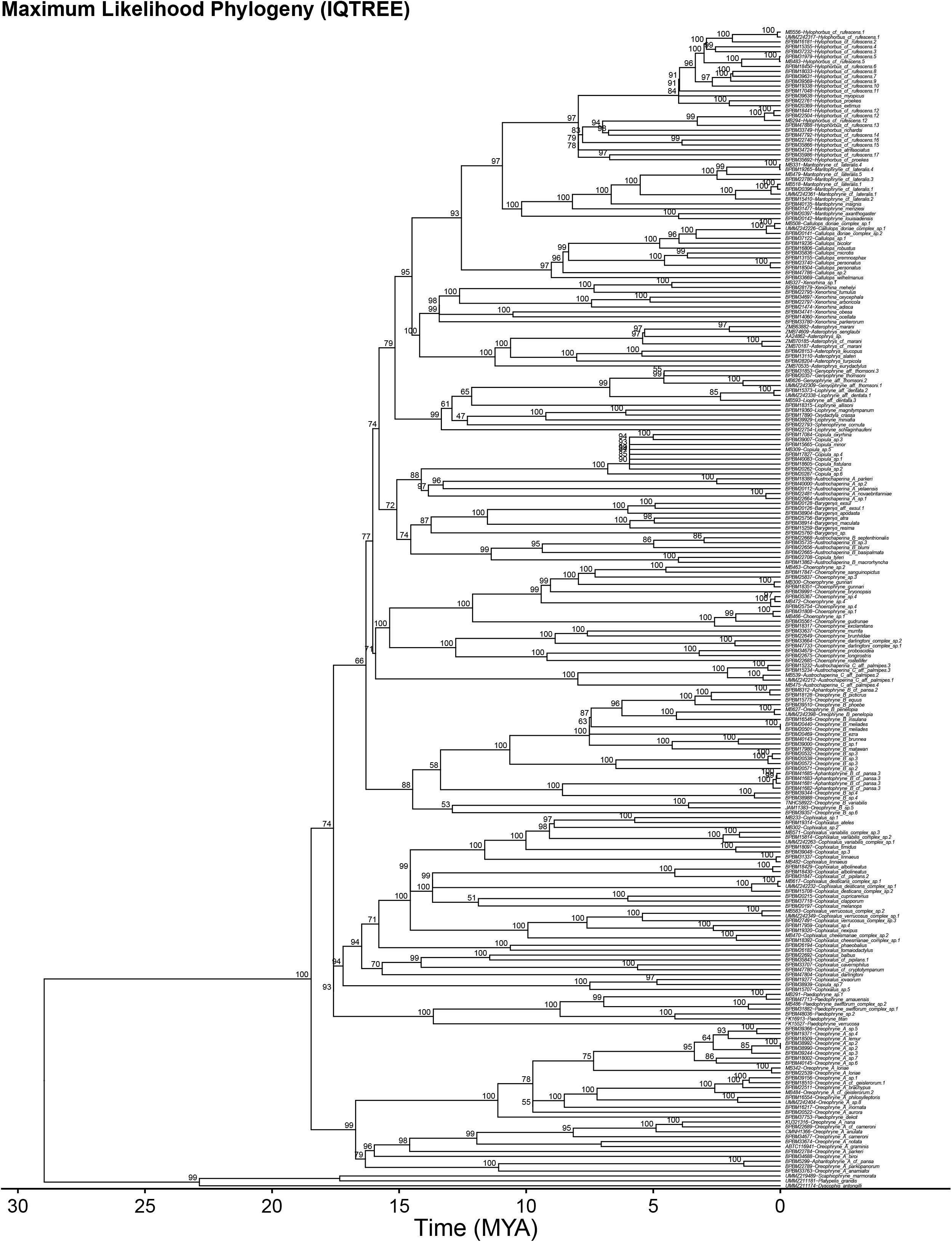
Time calibrated maximum likelihood phylogeny generated using IQTREE with bootstrap support indicated above nodes.

**Table 1:** Specimen metadata including collection site, type locality, species citation, lifestyle based on microhabitat use and citation, gps, elevation, site name and number, geological terrane, GenBank accession numbers by locus.

We also show nuclear-only (Figure 3) and mitochondrial-only (Figure 4) Bayesian inference trees, which are highly concordant in topology, although suffering a loss of resolution in relation to the nuclear+mitochondrial combined data tree. Therefore, the resolution of this phylogeny can only be achieved through the combination of nuclear and mitochondrial datasets (Figures 1-4). The nuclear-only tree recovered an age estimate for the clade of 30 MY, compared to 17 MY from the mitochondrial-only tree, and and age of 20 MY for the combined dataset, which are concordant with other estimates.

**Figure 3:**
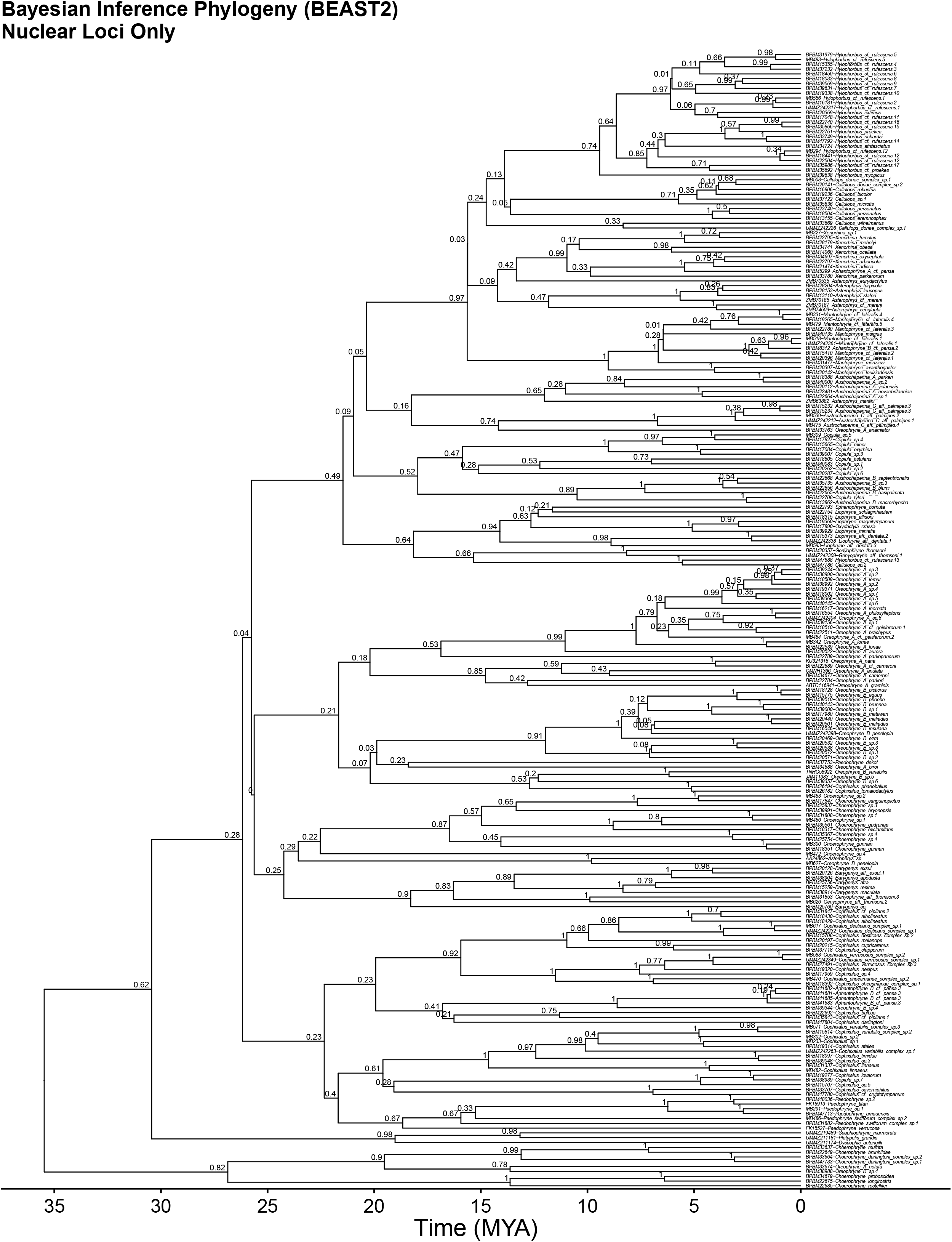
Nuclear-DNA-ooutgropnly time calibrated Bayesian inference phylogeny.

**Figure 4:**
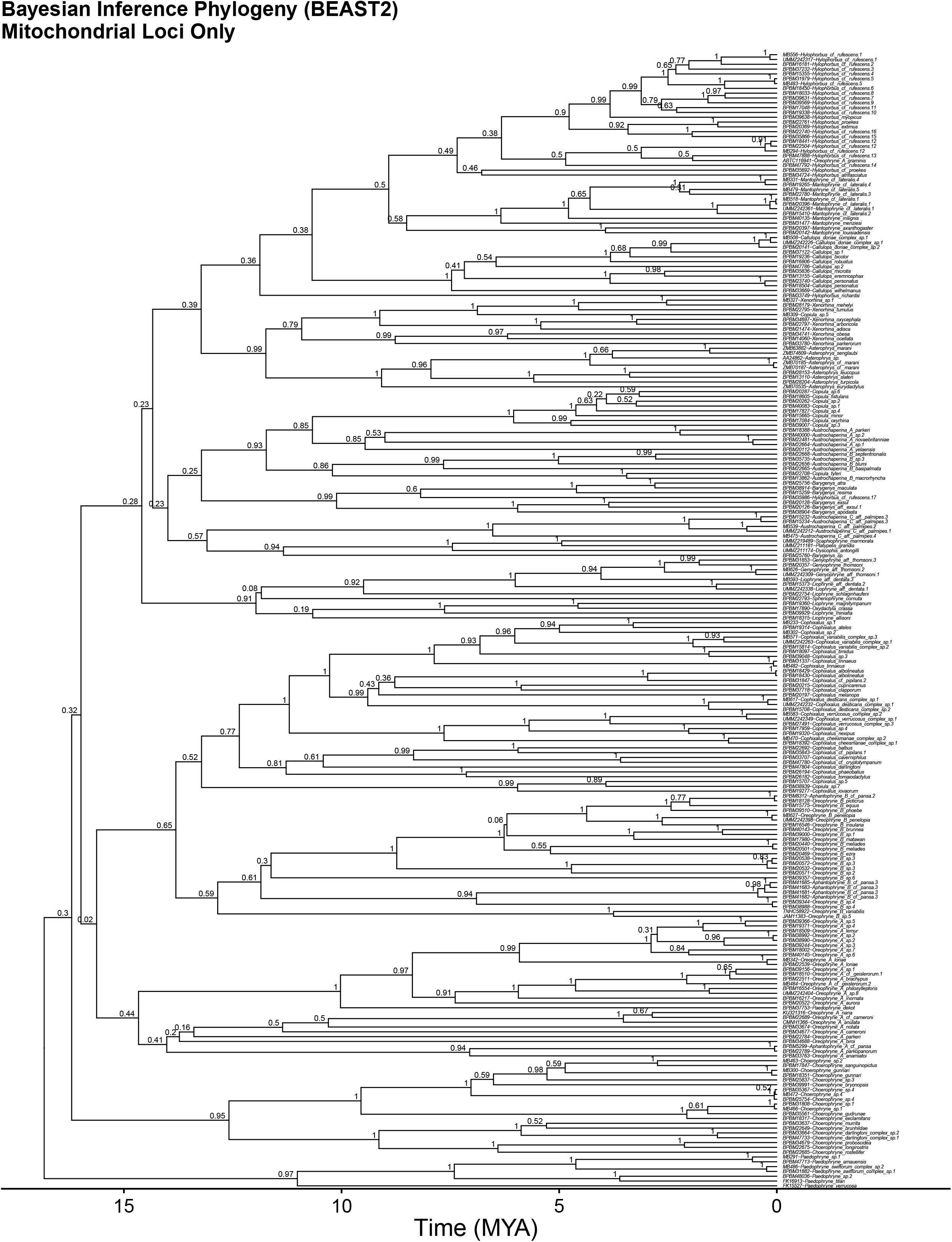
Mitochondrial-DNA-only time calibrated Bayesian inference phylogeny.

In addition, we present the specimen vouchers used in the phylogenetic analysis (Table 1), the species identification, collection site and locality, noting whether it was the type locality, species citation, GPS, elevation, site, geological terrane, lifestyle based on microhabitat use with their citations, and GenBank accession numbers by locus.

We developed new primers three nuclear (Seventh in Absentia (SIA), Brain Derived Neurotrophic Factor (BDNF), Sodium Calcium Exchange subunit-1 (NXC-1)), and two mitochondrial loci (Cytochrome oxidase b (CYTB), and NADH dehydrogenase subunit 4 (ND4)) loci (Table 2). We present the aligned dataset of 2474 base pairs (as input datasets for BEAST2 or IQTREE) that is 99% complete along with the data partitions, evolutionary models, and geological timings used to reconstruct the phylogeny ([1]; Figure 3; Table 3).

**Table 2:** Primer sequences redesigned for this study, based on Rivera et al. [15]. Product length reported in base pairs, and Tm in °C. Starting touchdown PCR temperature range in °C is reported in TD range. Outer indicates the outermost most forward/reverse primer for a locus.

Furthermore, we mapped anuran lifestyles (Table 1; one of arboreal, scansorial, terrestrial, fossorial, and semi-aquatic) onto the phylogeny (Figure 5), which indicated a general trend of niche conservatism throughout the clade [1]. Code to reproduce all analyses with all input files are provided in the GitHub repository.

**Figure 5:**
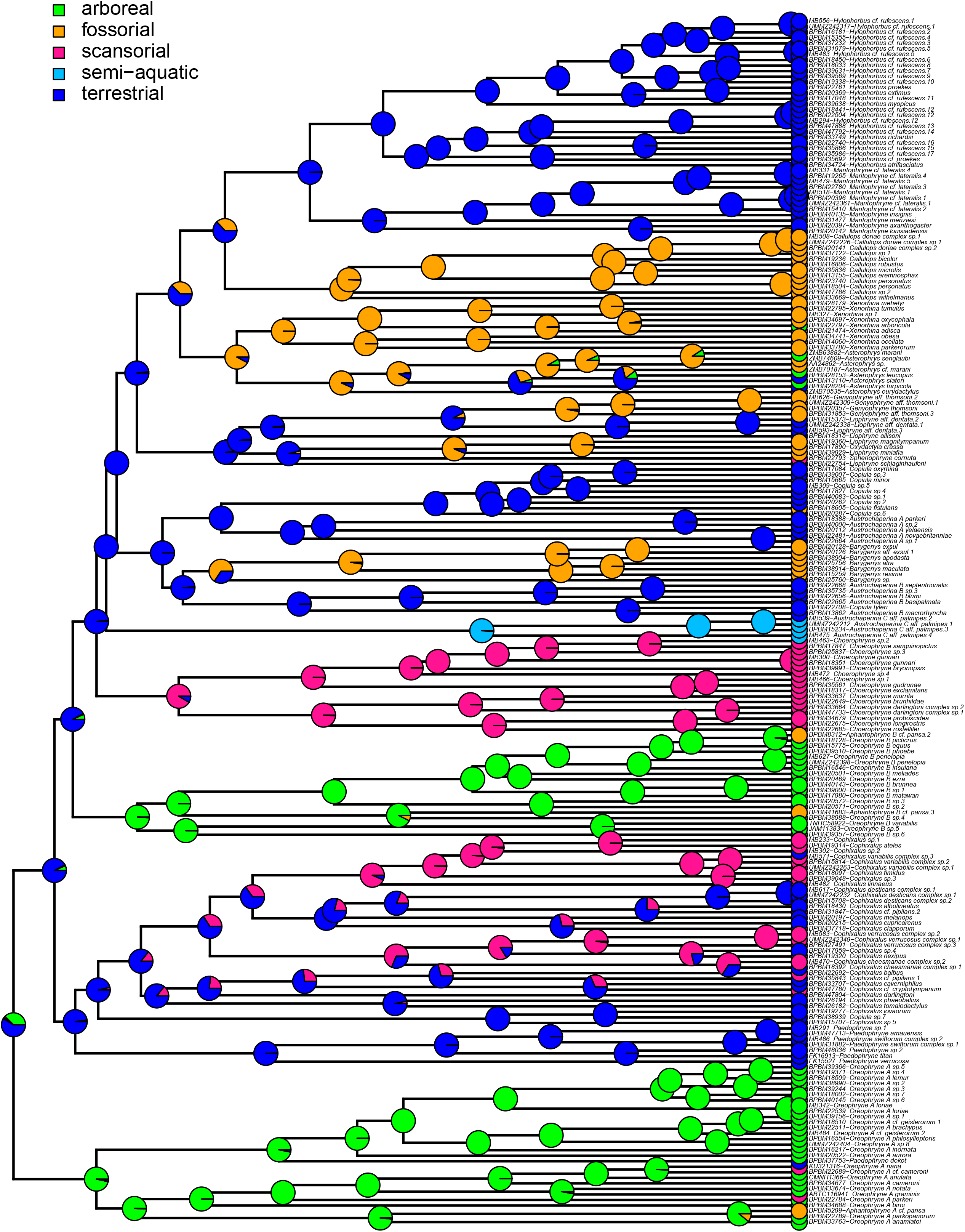
The evolution of Asterophryinae lifestyle. Lifestyles were mapped onto the 218 taxa time calibrated Bayesian inference phylogeny with ancestral states reconstructed using maximum likelihood. Pies on each node show the probabilities of lifestyles at ancestral nodes generated from 2000 stochastic mappings.

## Experimental design, materials and methods

### Sampling Design

We built upon a previous sequencing dataset [15] adding an additional 62 taxa to produce the largest dataset to date of 205 candidate taxa of Asterophryinae and three outgroup taxa [1]. We conducted new field collections of four New Guinea island sites and two offshore island sites (Normanby Island; site detail provided in Table 1).

Collections from the Central Highlands added samples to a region with low sampling. Furthermore, we included multiple samples of nominal species with “widespread distributions”, including at least one sample per site. We collected locality data from museum databases and identified 19 sites (18 multispecies) with reasonably complete species sampling.

### Primer Design

The previous dataset had significant missing data for some of the loci, in particular ND4 and NXC-1. We redesigned all primers (Table 2) for improved PCR performance in order to complete the dataset for all species. Nuclear primers were based on anuran and vertebrate sequences downloaded from GenBank [16] using NCBI Primer-BLAST software [2]. Mitochondrial sequences for Asterophryinae were downloaded from GenBank and aligned with previous sequencing efforts to design degenerate primers using HYDEN [17] with default settings.

### PCR Strategies to Improve Data Completeness

Standard PCR protocols were described in [1]. Several modifications were used to improve recovery of difficult targets. Touchdown PCR used annealing temperatures starting at 5 °C above Tm (Table 2) and decreasing by 1 °C per cycle for the first 10 cycles, followed by 25 cycles at the lowest annealing temperature. Some primer combinations for NXC-1, ND4, and CYTB required a touchdown window of 55-45 °C, whereas SIA and BDNF primers were improved by windows of 66-56 °C or 63-53 °C, respectively. Difficult targets were temperature optimized. A 15 degree touchdown window was used for CytB, as necessary, to improve yield likely due to the degeneracy or the greater Tm disparity between forward and reverse primers. Yield was improved for faint PCR bands with five additional amplification cycles. In other cases, a nested PCR strategy was effective with an initial PCR reaction using outer primers as the template for a nested or semi-nested reaction. PCR products were cleaned using Monarch spin columns (NEB). Several samples produced two bands for ND4, in these cases, bands of the correct length were excised from the gel and purified prior to Sanger sequencing. Sanger sequencing was conducted at the University of Hawaii at Manoa’s Advanced Studies of Genomics, Proteomics and Bioinformatics facility (https://www.hawaii.edu/microbiology/asgpb/).

### Increased Data Quality

We improved data quality by increasing data vetting beyond standard quality score criteria (>40%) and BLAST confirmation of loci. To eliminate the possibility of amplifying non-homologous targets such as pseudogenes, we furthermore instituted a practice of confirmation of suspect sequences by resequencing any data that resulted unexpected placement of taxa in single-locus phylogenies, sometimes with greater PCR stringency, as well as scrutinizing pairwise alignments of the data resulting in unexpected distance scores.

Sequence data were aligned after translation into amino acid sequences using MUSCLE [4], then back translated to nucleotides for phylogenetic reconstruction. Alignments provided in the Supplementary Materials.

### Phylogenetic Analysis

A more complex evolutionary model was used for phylogenetic analysis as compared to previous efforts. The best model was identified by simultaneously optimizing evolutionary models and partitions (by locus and codon) using PartitionFinder2 [5], and contained 13 partitions for the 5 locus dataset [(Table 3). Trees were time-calibrated using the timing of six geological events separating sister taxa (see [1] and file asterophryinae_dates.txt in Supplementary Materials). The maximum likelihood (ML) tree was estimated using IQTREE [7] with nodal support based on 2000 bootstrap replicates. The Bayesian inference (BI) tree was estimated using BEAST2 [6] with two independent Monte Carlo (MCMC) runs performed for 100,000,000 generations with sampling every 10,000 generations and 30% burn-in. Input files for these analyses are provided in Supplementary Materials. Further details of phylogenetic reconstruction are given in [1]. We checked for the possibility of morphologically cryptic species by examining mitochondrial divergence and phylogenetic distance [see 1].

**Table 3:** Data partitions and their best-fit evolutionary models for sequence evolution using PartitionFinder2 allowing models to vary by locus and codon position (15 possible partitions).

### Assembly of Metadata

Collection site data including GPS and elevation information were collected using museum databases for our voucher specimens as well as data collected from our own fieldwork. Samples were mapped to terranes (major geological sectors) using their GPS coordinates. Lifestyle data were collected based on microhabitat use (perch type) and behavior data from literature research and our own fieldwork. Briefly, frogs were classified as: arboreal if found on tree trunks or canopy, more than 2m above the ground; scansorial if found on shrubs less than 2m above the ground; terrestrial if generally found on the ground on the forest floor or among leaf litter; fossorial if found underground in holes or burrows; and semi-aquatic if found associated with streams and using swim for escape. All metadata, including museum number and locus-specific GenBank numbers are assembled in Table 1.

We evaluated the designation of a “collection site” empirically. Traditionally, field workers in New Guinea designate sites or “field camps” at various distances based on their perceptions of variation in the species under study. We analyzed spatial clustering of samples by calculating the distances between collection sites from their GPS coordinates using the geodist function in the R package geodist [10]. A spatial cutoff of 1.1 km corresponded well with collection sites designated by field workers, as well as with distinct species in our phylogeny (with the exception of cryptic species complexes). The sites are listed in Table 1.

### Ancestral Reconstruction of Lifestyle

We reconstructed ancestral states for lifestyle along the phylogeny using Maximum Likelihood (ML) with the fitDiscrete function in the R package geiger [8]. The best fit ML model assumed a symmetric transition rate between lifestyles (AICc=309) and was used to generate 2000 stochastic mappings of lifestyle onto the phylogeny using SIMMAP [18] with the make.simmap function in the R package phytools [9]. A majority of nodes (excepting 10) were reconstructed with >= 70% support for a single lifestyle (Figure 5).

Phylogenetic plots were created using the R packages ggtree, ouch and treeio [11, 12, 13], and with a custom script based on modifications of ouch. All analyses were conducted in the R statistical computing environment unless otherwise noted [14]. Code for all analyses are provided in the supplementary file “asterophryinae_phylogeny_analysis_figures.R”. All supplementary materials are available in the repository (https://doi.org/10.5281/zenodo.7063167).

## Ethics statements

No human subjects were used in this study. Animals used in this study were euthanized using MS-222 following UH IACUC protocol 12–1458 issued to M. Butler.

## CRediT author statement

**Ethan C. Hill:** Formal analysis, Data Curation, Methodology, Project administration, Writing - Original Draft, Investigation, Writing - Review & Editing, Visualization **Claire J. Fraser:** Investigation, Data Curation, Writing - Original Draft, Writing - Review & Editing **Diana F. Gao:** Investigation, Writing - Review & Editing, Visualization **Mary J. Jarman:** Investigation, Writing - Review & Editing **Elizabeth R. Henry:** Investigation, Resources, Data Curation **Bulisa Iova:** Investigation, Resources **Allen Allison:** Conceptualization, Methodology, Investigation, Resources, Supervision, Writing - Original Draft, Writing - Review & Editing **Marguerite A. Butler:** Conceptualization, Methodology, Formal analysis, Data Curation, Supervision, Investigation, Writing - Original Draft, Writing - Review & Editing, Visualization, Project administration, Funding acquisition, Visualization

## Funding

This study was made possible by a grant from the National Science Foundation awarded to MB (DEB-1145733). Some of the specimens were jointly collected with Fred Kraus (DEB-1345063).

## Specimen Loans

Specimens were kindly provided by Molly Hagemann and Pumehana Imada (Bishop Museum, Honolulu), Rainer Günther (Museum für Naturkunde, Berlin), Jim McGuire (Museum of Vertebrate Biology, Berkeley), Rafe Brown (University of Kansas, Lawrence), and Fred Kraus, Ron Nussbaum, and Gregory Schneider (University of Michigan Museum of Zoology, Ann Arbor).

We thank Fred Kraus and Julio Rivera for consultation early in this work. Julio Rivera, Jeff Scales, Nalani Kito-Ho, Niegel Rozet, Jeff Higa and Raine Higa provided generous assistance in the field. We are grateful to the field assistance from many local residents: at Maru Ruama, Mt. Geregu: Peter Joseph, Dambio Moi, Walter Moi, David Peter, and Laiwoi Yoiini; at Cliffside Camp, Kamiali: Maties Dagam, David Enoch, Lenny Keputong, and Marcus Symon; at Normanby Island: Clement Bobby, Fred Francisco, Kenny Lakson, Waiyaki Nemani, James Waraia, and Roland Waraia, and Misima Island: Simon Emidi and Kelly Nabwakulea. We thank Normanby Mining PNG, LTD for land access and accommodation. We thank Georgia Kaipu of the PNG National Research Institute) and the late Barnabus Wilmont of the PNG Department of Environment and Conservation for assistance with research permits and necessary visas, and Andrew Moutu, the Director of the PNG National Museum and Art Gallery, for use of facilities.

We are grateful for excellent critical comments from Alex Slavenko, Paul Oliver, Dave Carlon, and three anonymous reviewers that greatly improved the manuscript.

## Declaration of interests

☒ The authors declare that they have no known competing financial interests or personal relationships that could have appeared to influence the work reported in this paper.
□ The authors declare the following financial interests/personal relationships which may be considered as potential competing interests:

## Notes

https://doi.org/10.5281/zenodo.7063167

### Competing Interest Statement

The authors have declared no competing interest.

